# Thresholds of drought and terrain complexity shape biomass patterns in South America’s Caatinga

**DOI:** 10.1101/2025.11.02.684225

**Authors:** Brhenda Santos Lozado, Cinthia Pereira de Oliveira, Alessandro de Paula, Patrícia A. B. Barreto-Garcia, Odair Lacerda Lemos, Ana Luisa Leite Pereira, Emanuel Araújo Silva, Rinaldo L. C. Ferreira, José A. Aleixo da Silva, Frans Pareyn, Peter W. Moonlight, Domingos Cardoso, Elmar Veenendaal, Luciano Paganucci de Queiroz, Priscyla M. S. Rodrigues, Rubens Manoel dos Santos, Tiina Sarkinen, Toby Pennington, Oliver L. Phillips, Robson Borges de Lima

**Affiliations:** Universidade Estadual do Sudoeste da Bahia; Universidade do Estado do Amapá; Universidade Federal de Santa Maria; Universidade Federal Rural de Pernambuco; Associação Plantas do Nordeste; Royal Botanic Garden Edinburgh; Universidade Federal da Bahia; Wageningen University; Universidade Estadual de Feira de Santana; Universidade Federal do Vale do São Francisco; Universidade Federal de Lavras; University of Exeter; University of Leeds

**Author notes:** Correspondence author: rbl.

**Keywords:** tropical dry forests, spatial modeling, hydroclimatic stress, climatic resources, forest inventory, global carbon cycle, climate change mitigation

## Abstract

Seasonally Tropical Dry Forests (STDFs) are widespread throughout the world and store significant amounts of carbon; however, they are often overlooked in spatial assessments compared to humid tropical forests. The Caatinga, which is the largest seasonally dry tropical forest in South America, covers approximately 862,000 km^²^ in northeastern Brazil and supports millions of people. Unfortunately, its carbon dynamics has not been thoroughly quantified, especially after centuries of land-use transformation and wood extraction that have significantly diminished its biomass stocks. In this study, we model the potential aboveground biomass (AGB) that Caatinga could sustain under current climatic, atmospheric, and topographic conditions. We integrated data from 301 geo-referenced plots along with high-resolution environmental predictors. The principal Component Analysis revealed two main gradients: a hydro-thermal axis dominated by precipitation, temperature, and severity of drought, and a topographic axis reflecting slope, ruggedness, and terrain position. Random Forest models, validated through both random and spatial cross-validation, explained a significant amount of variation in AGB (R^²^ = 0.81, RMSE = 21 Mg ha). Our findings indicated that AGB is highly sensitive to water availability. In particular, biomass increased sharply when the maximum cumulative water deficit (MCWD) was less than –500 mm and annual rainfall exceeded approximately 1,100 mm. In contrast, elevated vapour pressure deficit (VPD) and potential evapotranspiration (PET) were associated with reduced carbon storage. Topographic heterogeneity further influenced AGB, with rugged and concave terrains supporting potential biomass levels more than twice as high as those found in flat, convex areas. Our predictive map reveals a mosaic of low-biomass cores and localized high-biomass refugia, highlighting the dual influence of hydroclimatic and topographic factors. These findings reposition Caatinga as a heterogeneous and dynamic potential carbon reservoir – historically degraded by changes in land-use but still capable of storing substantial carbon. This research offers critical insights for the restoration, conservation and mitigation efforts for climate change in tropical drylands worldwide.

## 1 Introduction

Tropical forests play a central role in the global carbon cycle, acting simultaneously as vast reservoirs and dynamic sinks of atmospheric carbon dioxide. While humid tropical forests, particularly the Amazon, Congo, and Southeast Asian rainforests, have been extensively studied for their carbon storage and fluxes (Baccini & Asner, 2013; Ferraz et al., 2018; Lewis et al., 2013; S. Saatchi et al., 2009), Seasonally Tropical Dry Forests (STDFs) remain comparatively overlooked. This imbalance is striking given that STDFs cover nearly half of tropical forested areas worldwide (Da Silva et al., 2017; Miles et al., 2006; Sampaio et al., 2010) and provide essential ecosystem services to millions of people. Their ability to act as net carbon sinks is increasingly recognized, yet their vulnerability to climate extremes suggests that they may also switch to sources under unfavorable conditions (Anderson et al., 2018; Powers et al., 2018).

The Caatinga, the largest continuous seasonally dry tropical forest in the Americas, is a critical case for understanding biomass dynamics in water-limited environments. This biome occupies 862,000 km^2^ in northeastern Brazil, harbors high levels of endemism, and sustains the livelihoods of nearly 27 million people (Araujo et al., 2023; Da Silva et al., 2017). Despite its ecological and socioeconomic importance, the Caatinga remains underrepresented in global assessments of carbon dynamics (Althoff et al., 2018). Most spatial analyses of aboveground biomass (AGB) have concentrated on humid forests, leading to an incomplete picture of carbon storage across tropical biomes (Chave et al., 2014). This knowledge gap is particularly concerning given the intensification of droughts, rising temperatures, and land-use pressures in the semi-arid tropics.

Environmental gradients strongly shape spatial variation in biomass within STDFs, yet few studies have systematically quantified these relationships at the biome scale (Alves Rodrigues Pinheiro et al., 2016; Araujo et al., 2023; Cavalcante et al., 2023). In the Caatinga, the interplay of climatic factors (precipitation, temperature), atmospheric stressors (vapor pressure deficit, potential evapotranspiration, aridity, drought severity), and topographic heterogeneity (slope, ruggedness, elevation) is expected to modulate vegetation structure in complex ways (Campos et al., 2019; Mendes et al., 2020). Climatic resources such as precipitation promote biomass accumulation by alleviating water limitation, whereas atmospheric stressors exacerbate plant water loss, reduce stomatal conductance, and increase hydraulic failure risk (Apgaua et al., 2022; Bauman et al., 2022). At finer scales, topography redistributes soil water and creates microclimatic refugia that buffer plants against regional climatic stress (Siyum, 2020).

The physiological responses of woody plants to these gradients further underscore the importance of integrating multiple drivers. In water-limited systems, hydraulic traits such as xylem vulnerability to cavitation, rooting depth, and drought deciduousness directly influence biomass accumulation and persistence (Alves Rodrigues Pinheiro et al., 2016; Smith-Martin et al., 2023). Elevated vapor pressure deficit (VPD) and prolonged maximum cumulative water deficit (MCWD) increase transpirational demand and push plants toward hydraulic failure, while higher precipitation and soil moisture availability sustain growth and regeneration (Barkhordarian et al., 2019; Costa et al., 2017; Gauthey & Gardner, 2024; Schönbeck et al., 2022; Xie et al., 2025). Rugged terrain, by contrast, can provide critical microsites where woody plants accumulate greater biomass due to improved water retention and reduced evaporative stress (Mattos et al., 2023; Terra et al., 2018). Such mechanistic links highlight how the additive and interactive effects of hydroclimatic resources, atmospheric demand, and terrain complexity likely govern AGB distribution in the Caatinga.

Despite these well-founded ecological mechanisms, spatially explicit evaluations of Caatinga biomass remain scarce. Most available studies focus on local inventories or experimental plots (DRYFLOR et al., 2016), limiting the ability to generalize patterns across the biome. This lack of comprehensive spatial models hampers our capacity to anticipate how climate change, land-use intensification, and prolonged droughts may alter carbon storage. Filling this gap is urgent, as dry forests are projected to face disproportionately stronger climate extremes compared to humid forests, with implications for biodiversity, livelihoods, and global carbon budgets (K. Allen et al., 2017).

Global efforts have sought to map aboveground biomass at continental and planetary scales through satellite-driven models such as the ESA Climate Change Initiative (CCI), NASA’s GEDI LiDAR mission, and pan-tropical biomass maps by Baccini et al. (2017) and Saatchi et al. (2011). Although these products provide valuable benchmarks for understanding global carbon distribution, their coarse resolution (≈500 m–1 km) and limited calibration in seasonally dry ecosystems restrict their accuracy in highly heterogeneous biomes such as the Caatinga. At local and regional scales, promising advances have been made using airborne LiDAR and geostatistical modeling (Oliveira et al., 2021; Viana Santos et al., 2023), yet these remain localized efforts that capture only fragments of the biome’s environmental and structural diversity. Together, these limitations underscore the critical need for high-quality, in situ calibration datasets to anchor remote-sensing estimates to ecological reality.

Accurate biomass modeling in the Caatinga depends on carefully measured ground data that include tree-level allometry, species identification, and wood density–key parameters that allow the translation of remotely sensed canopy or volume proxies into actual biomass and carbon stocks. Such plot-based data provide the ecological foundation required to refine regional models and reduce uncertainty in carbon accounting (DRYFLOR et al., 2016; ForestPlots.net et al., 2021). By integrating this field-based information with high-resolution environmental predictors and machine-learning approaches, our study offers the first biome-wide model of potential aboveground biomass (AGB) for the Caatinga.

It is important to note that this model represents potential, rather than current, biomass distribution. Centuries of human occupation, wood extraction, grazing, and agricultural expansion have markedly reduced actual biomass across vast portions of the biome, lowering stocks well below their ecological capacity (Araujo et al., 2023). The field plots used here were established in relatively mature and minimally disturbed stands, thus approximating the upper envelope of biomass achievable under present climatic and topographic conditions. As such, the resulting spatial model reflects the biophysical potential of the Caatinga to store carbon in the absence of significant anthropogenic degradation, providing a valuable baseline for restoration planning, conservation prioritization, and assessment of deviations between current and potential carbon stocks.

Here, we hypothesize that biomass patterns in the Caatinga are primarily governed by water availability and stress gradients, with higher AGB in areas of greater precipitation and less severe drought (MCWD), and lower AGB under elevated atmospheric demand (VPD, PET) and extreme aridity. We further hypothesize that topographic complexity acts as an independent modulator, generating localized biomass refugia in rugged and concave terrains. At the same time, convex and exposed sites sustain lower stocks due to intensified evaporative stress. Collectively, these hypotheses posit that both large-scale hydroclimatic drivers and fine-scale terrain heterogeneity shape the Caatinga’s biomass. The objectives of this study are threefold: (i) to quantify and evaluate how climatic, atmospheric, and topographic factors interact to shape AGB distribution across the Caatinga; (ii) to generate a robust spatial model of AGB using Random Forests validated through random and spatial cross-validation; and (iii) to produce the first biome-wide predictive map of Caatinga potential biomass, with implications for conservation, restoration, and climate policy. By elucidating the environmental mechanisms underlying carbon storage in the world’s largest tropical dry forest, this study provides critical insights into the resilience and vulnerability of dryland ecosystems under global climate change.

## 2 Material and Methods

### 2.1 Study area

This study was conducted in the Caatinga, the largest seasonally dry tropical forest in the Americas and a global biodiversity hotspot for dryland ecosystems. The biome spans approximately 862,818 km^2^, covering ≈18 % of Brazil’s territory and sustaining the livelihoods of nearly 27 million people (IBGE, 2019). Its distribution extends across nine northeastern states–Alagoas, Bahia, Ceará, Maranhão, Pernambuco, Paraíba, Rio Grande do Norte, Piauí, and Sergipe–as well as the northern portion of Minas Gerais (Figure 1).

**Figure 1:**
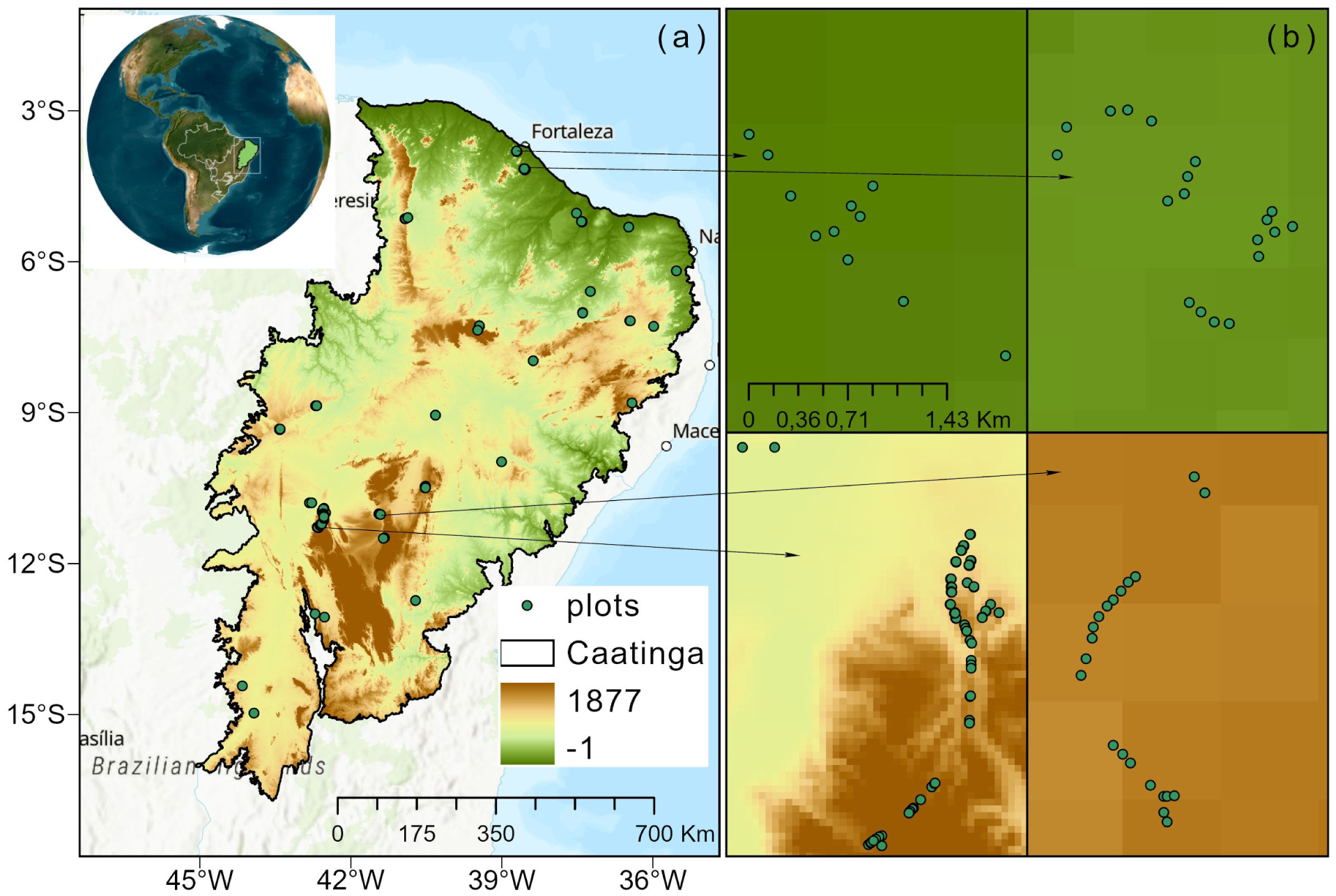
Study area and distribution of field plots across the Caatinga biome. (a) Geographic extent of the Caatinga, the largest seasonally dry tropical forest in the Americas, spanning ≈862,818 km^2^ in northeastern Brazil. The biome’s location within South America is shown in the inset. Green circles indicate the 301 geo-referenced field plots distributed across 31 sites, used to estimate aboveground woody biomass. The color gradient represents terrain elevation above sea level in metres, highlighting the firm environmental heterogeneity of the region. (b) Enlarged views of four quadrants illustrate clusters of plots capturing contrasting topographic conditions, from lowland depressions to higher-elevation plateaus. These field sites encompass gradients of altitude and terrain roughness that shape vegetation structure and biomass storage

The regional climate is classified as hot semi-arid (BSwh, Köppen system), with mean annual temperatures ranging from 25 °C to 30 °C and annual rainfall between 400 and 1200 mm. Rainfall is markedly irregular, concentrated within three to four months, while the dry season may last 7–9 months, one of the most distinctive climatic features of the biome (Da Silva et al., 2017). Soils are typically shallow, rocky, and prone to salinization due to low precipitation and high evapotranspiration rates. The dominant soil groups include Argisols, Latosols, and Luvisols, which strongly constrain vegetation structure and primary productivity (Santos, 2018).

Vegetation in the Caatinga is classified as wooded steppe-savanna, encompassing a mosaic of woody species, herbaceous layers, cacti, and bromeliads. These plant communities exhibit marked adaptations to water scarcity and thermal stress, including deciduousness, succulence, and specialized root systems. The flora comprises at least 3,150 species distributed across 950 genera, with ≈23% endemism, including 29 genera restricted to the biome (Amorim et al., 2005; DRYFLOR et al., 2016). This combination of climatic harshness, edaphic constraints, and evolutionary adaptation makes the Caatinga a unique natural laboratory for assessing biomass dynamics in seasonally dry tropical systems.

### 2.2 Biomass database

The aboveground biomass (AGB) dataset was compiled from 301 geo-referenced plots and distributed across 31 sites spanning the Caatinga biome (Figure 1). These plots were established in forest fragments representing the broad ecological spectrum of Caatinga vegetation, with inventories sourced from Instituto do Meio Ambiente e Recursos Hídricos-Inema, the Rede de Manejo Florestal da Caatinga (RMFC), and ForestPlots.net (ForestPlots.net et al., 2021). Together, these sources provide a comprehensive sampling of vegetation structure across gradients of climate, soils, and land-use history in northeastern Brazil.

Plot dimensions varied according to survey protocols and institutional objectives. While most followed the standard size of 20 × 20 m (400 m^2^), others were 10 × 20 m (200 m^2^), 20 × 30 m (600 m^2^), 15 × 50 m (750 m^2^), 20 × 50 m (1,000 m^2^), and 100 × 100 m (1 ha). This heterogeneity in sampling design was harmonized by standardizing the minimum inclusion criterion to trees with diameter at breast height (DBH) ≥ 3 cm, measured at 1.30 m above ground level. This criterion ensured comparability among inventories while capturing the majority of aboveground biomass stored in woody vegetation.

AGB for each tree was estimated using the allometric model developed by Sampaio et al. (2010), calibrated explicitly for dry forests of northeastern Brazil and widely applied in Caatinga biomass assessments. The equation is expressed as:

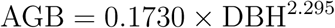

**Where:** AGB represents the individual aboveground dry biomass (kg), and DBH the trunk diameter (cm). This model has demonstrated strong predictive performance (*R*^2^ = 0.92), capturing the nonlinear scaling between stem diameter and biomass accumulation in semi-arid tropical forests. Tree-level AGB estimates were subsequently aggregated to the plot level and standardized to Mg ha*^−^*^1^, enabling spatial comparisons across sites with variable plot sizes and survey designs.

### 2.3 Environmental predictors

To investigate the environmental drivers of aboveground biomass (AGB) variation across the Caatinga, we compiled a suite of climatic, atmospheric, and topographic variables widely recognized as critical determinants of vegetation structure in seasonally dry tropical forests. These predictors represent either resource-related factors that enhance biomass accumulation (e.g. water availability, precipitation), or stressors/disturbance-related factors that limit biomass through physiological stress or environmental constraints (e.g. vapor pressure deficit, aridity, slope exposure) (Table 1).

**Table 1:**
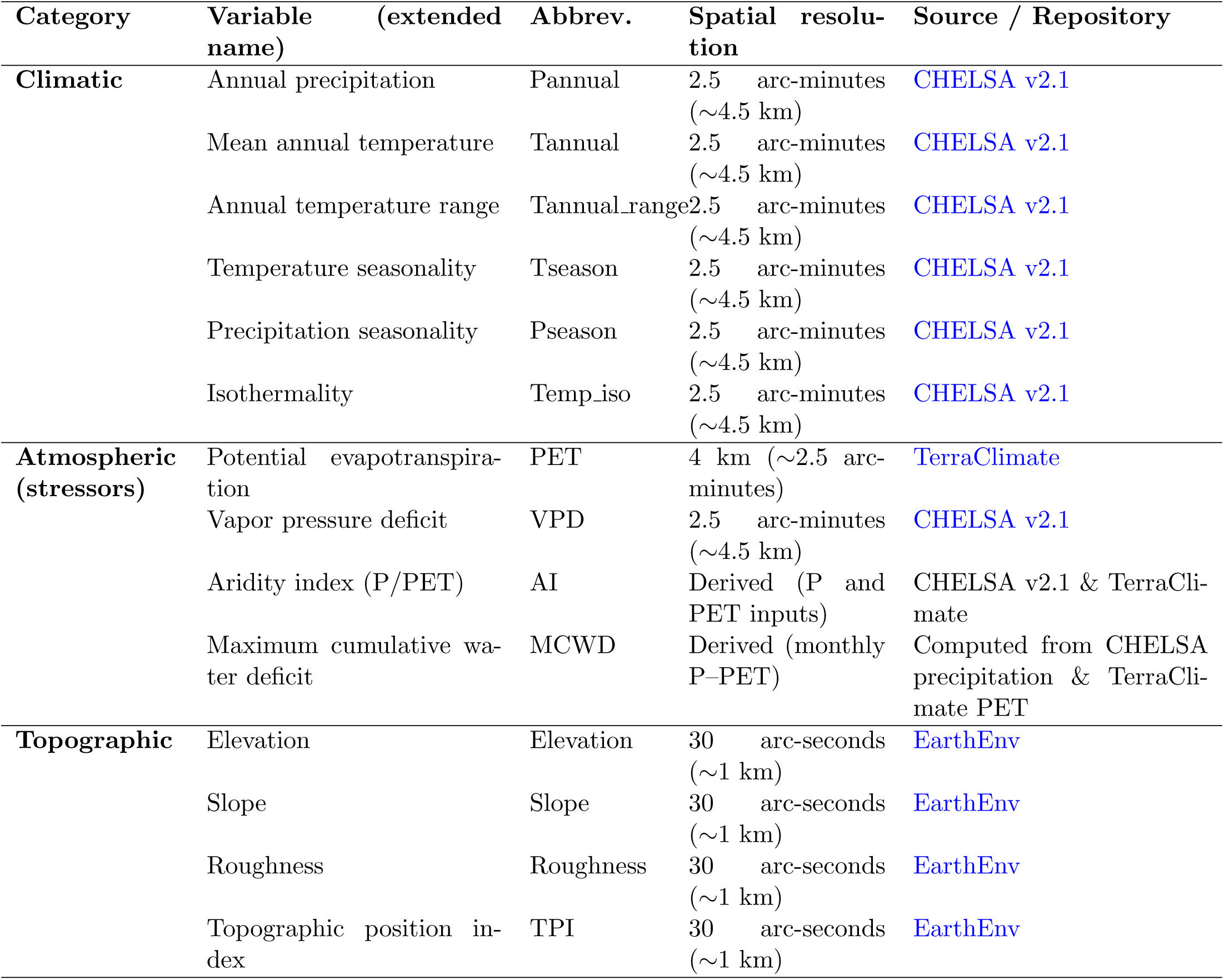
Environmental predictors used to model aboveground biomass (AGB) across the Caatinga. Climatic, atmospheric, and topographic predictors were assembled from global repositories to represent resource availability (e.g., precipitation, temperature) and environmental stressors (e.g., vapor pressure deficit, aridity, MCWD) influencing biomass accumulation in seasonally dry tropical forests. All predictors were standardized to a common resolution of 30 arc-seconds (∼1 km) and cropped to the Caatinga domain. Climatic variables were obtained from CHELSA v2.1, atmospheric stressors from CHELSA and TerraClimate, and topographic metrics from EarthEnv. The table details the extended variable name, abbreviation, spatial resolution, and source repository with active links.

| Category | Variable (extended name) | Abbrev. | Spatial resolution | Source / Repository |
| --- | --- | --- | --- | --- |
| **Climatic** | Annual precipitation | Pannual | 2.5 arc-minutes ($\sim 4.5$ km) | CHELSA v2.1 |
| | Mean annual temperature | Tannual | 2.5 arc-minutes ($\sim 4.5$ km) | CHELSA v2.1 |
| | Annual temperature range | Tannual_range | 2.5 arc-minutes ($\sim 4.5$ km) | CHELSA v2.1 |
| | Temperature seasonality | Tseason | 2.5 arc-minutes ($\sim 4.5$ km) | CHELSA v2.1 |
| | Precipitation seasonality | Pseason | 2.5 arc-minutes ($\sim 4.5$ km) | CHELSA v2.1 |
| | Isothermality | Temp_iso | 2.5 arc-minutes ($\sim 4.5$ km) | CHELSA v2.1 |
| **Atmospheric (stressors)** | Potential evapotranspiration | PET | 4 km ($\sim 2.5$ arc-minutes) | TerraClimate |
| | Vapor pressure deficit | VPD | 2.5 arc-minutes ($\sim 4.5$ km) | CHELSA v2.1 |
|  | Aridity index (P/PET) | AI | Derived (P and PET inputs) | CHELSA v2.1 & TerraClimate |
|  | Maximum cumulative water deficit | MCWD | Derived (monthly P–PET) | Computed from CHELSA precipitation & TerraClimate PET |
| **Topographic** | Elevation | Elevation | 30 arc-seconds ($\sim 1$ km) | EarthEnv |
| | Slope | Slope | 30 arc-seconds ($\sim 1$ km) | EarthEnv |
| | Roughness | Roughness | 30 arc-seconds ($\sim 1$ km) | EarthEnv |
| | Topographic position index | TPI | 30 arc-seconds ($\sim 1$ km) | EarthEnv |

Climatic variables included annual precipitation (P_annual_), temperature seasonality (T_season_), precipitation seasonality (P_season_), annual temperature range (T_annual range_), mean annual temperature (T_annual_), and isothermality (Temp_iso_). Precipitation provides direct water input to support productivity, while seasonal and interannual variability impose constraints on vegetation persistence. These data were obtained from the CHELSA v2.1 repository at 2.5 arc-minute spatial resolution (Karger et al., 2017).

Atmospheric variables captured the role of evaporative demand and drought severity. Potential evapotranspiration (PET) was obtained from TerraClimate at ∼4 km spatial resolution (Abatzoglou et al., 2018), while vapor pressure deficit (VPD) was derived from CHELSA (2.5 arc-minutes). Both variables represent atmospheric stressors: higher PET and VPD increase transpirational costs and risk of hydraulic failure, negatively impacting biomass accumulation (Bauman et al., 2022; Mattos et al., 2023). The aridity index (AI) was derived as the ratio of precipitation to PET (AI = P*/*PET), representing long-term water balance and plant-available moisture (Zomer et al., 2022).

We further incorporated the Maximum Cumulative Water Deficit (MCWD), an integrative measure of drought severity widely used in tropical forest ecology (Campanharo & Silva-Junior, 2019; Xie et al., 2025). MCWD corresponds to the most negative cumulative water balance (precipitation minus PET) reached within a given year at each pixel, thus quantifying the depth of seasonal drought. Monthly deficits (P_annual_ − PET) were accumulated sequentially, with the lowest annual value recorded as the MCWD. This variable provides a sensitive indicator of prolonged dry-season stress, strongly associated with tree mortality and biomass decline across tropical regions (Xie et al., 2025).

Topographic variables were derived from the EarthEnv topographic database (Amatulli et al., 2018) at 30 arc-seconds (∼1 km). These included elevation, slope, roughness, and topographic position index (TPI). Topography influences soil depth, drainage, microclimate, and resource redistribution, creating local heterogeneity that may either buffer climatic stress (e.g., valleys, concave slopes) or exacerbate it (e.g., ridges, convex landforms).

All predictors were cropped to the Caatinga domain and re-sampled to a common spatial resolution of 30 arc-seconds (∼1 km) to ensure comparability across layers and integration into the spatial modeling framework. Subsequent processing was conducted in R v4.4.2 (R Core Team, 2024), where geospatial covariates were extracted at the coordinates of each sampling plot using the raster::extract function from the raster package (Hijmans et al., 2021). This procedure ensured consistent retrieval of environmental values at the plot or fragment level, which were then organized into a predictor matrix and used for statistical analyses and model fitting.

### 2.4 Data analysis and spatial modeling

To explore how environmental gradients structure aboveground biomass (AGB) across the Caatinga, we employed a combination of multivariate ordination and machine learning approaches, allowing both visualization of environmental drivers and predictive modeling of biomass distribution.

First, we performed a Principal Component Analysis (PCA) to summarize the multidimensional structure of the environmental space. PCA was applied using the FactoMineR package (Husson et al., 2025) to standardized climatic, atmospheric, and topographic predictors to identify dominant axes of variation and evaluate how they align with biomass distribution. This ordination provided a synthetic view of the hydro-thermal continuum (precipitation, temperature, and water deficit) versus terrain complexity (slope, ruggedness, and topographic position), highlighting the degree of independence between large-scale climatic gradients and local topographic modulators. The PCA biplot was used as an exploratory tool to visualize how combinations of environmental drivers potentially constrain or promote AGB.

For predictive modeling, we implemented Random Forests (RF) using the randomForest package (Cutler & Wiener, 2022), a non-parametric ensemble method widely applied in ecological modeling due to its ability to capture non-linear relationships and complex interactions among predictors (Calhoun et al., 2020; Genuer & Poggi, 2020). RF models were trained using tree-level AGB aggregated to plot-level values (Mg ha*^−^*^1^) as the response variable, with environmental predictors as covariates.

We evaluated model performance using two complementary strategies: (i) random *k*-fold crossvalidation, partitioning the dataset into 15 folds to assess predictive accuracy under random splits, and (ii) to minimize potential biases arising from spatial autocorrelation, which can artificially inflate model accuracy due to the proximity of sampling plots, we implemented a spatial cross-validation framework. Instead of random partitioning of the dataset, which tends to assign geographically close plots to both training and testing sets, the spatial cross-validation grouped sites into spatially disjoint folds based on their geographic coordinates. This procedure ensured that training and validation data were separated by explicit spatial distance, reducing the effect of local neighborhood similarity and providing a more conservative and robust estimate of model performance.

By adopting this strategy, the evaluation metrics (*R*^2^ and RMSE) better reflect the model’s capacity to generalize across the full environmental and geographic heterogeneity of the Caatinga, rather than being influenced by localized clustering of plots. Such an approach has been increasingly recommended in ecological modeling studies (Roberts et al., 2017; Valavi et al., 2018; Liang et al., 2022), as it accounts for the inherent spatial structure of biodiversity and biomass datasets and provides more reliable inferences for large-scale predictions.

Model accuracy was assessed through multiple metrics: the coefficient of determination (*R*^2^), root mean squared error (RMSE), and mean absolute error (MAE), complemented by calibration plots to evaluate systematic bias. To interpret model outcomes, we examined partial dependence plots (PDPs), which reveal the marginal effect of each predictor on AGB while holding others constant, thereby identifying nonlinear responses and ecological thresholds (Cutler & Wiener, 2022). In parallel, we quantified predictor importance using permutation-based measures of mean decrease in accuracy and node purity (IncMSE, IncNodePurity), highlighting the relative influence of climatic versus topographic controls.

Following model training and validation, the fitted Random Forest was applied in a spatial prediction framework in which each pixel of the coregistered environmental covariates was passed through the ensemble of decision trees. Rather than relying on explicit coefficients or intercepts, as in parametric regression models, the Random Forest generates predictions by aggregating the outputs of multiple recursive partitioning trees through a majority-vote or mean-ensemble approach. Each pixel was thus classified according to the hierarchical splits defined by the predictor variables, and its predicted value represented the average across the ensemble of trees. By propagating this process across the full stack of raster layers, we produced a continuous biome-wide map of aboveground biomass (AGB) at ∼1 km spatial resolution, directly linking the environmental gradients of climate, atmospheric demand, and topography to spatially explicit estimates of carbon storage.

To assess the degree of overlap and independence among the key environmental drivers influencing aboveground biomass (AGB) at the plot level, we performed an UpSet intersection analysis (Lex et al., 2014). The five predictors used in this analysis were selected based on their high relative importance scores in the Random Forest model. Each variable was categorized into binary classes representing extreme environmental conditions. The UpSet framework quantifies how often these extreme conditions occur individually or in combination across the 301 sampling plots, thereby revealing whether single stressors or synergistic interactions drive local biomass variation among multiple factors. This approach provides a robust visualization of the intersection structure among the most influential environmental gradients, clarifying how additive versus interactive effects shape spatial patterns of AGB in the Caatinga.

## 3 Results

### 3.1 Regional patterns of AGB

The controls of aboveground biomass (AGB) in the Caatinga emerge along two nearly orthogonal axes of variation (Figure 2). In the PCA biplot, the first component (Dim1, 39.2% of variance) reflects a hydro-thermal continuum. Annual precipitation (up to ∼1,200 mm), mean annual temperature, and isothermality align with atmospheric demand metrics such as vapor pressure deficit (VPD), potential evapotranspiration (PET), and maximum cumulative water deficit (MCWD). Higher precipitation and less negative MCWD values (−700 to −500 mm) cluster on the right side, while stronger temperature seasonality projects to the left, opposing this climatic gradient. The second component (Dim2, 21.0% of variance) separates topographic attributes–slope, roughness, and elevation–which load strongly above the horizontal axis and remain distinct from climatic drivers. This orthogonality indicates that relief effects operate as independent local modulators of water redistribution and microclimate, with potential to amplify or buffer regional climatic constraints.

**Figure 2:**
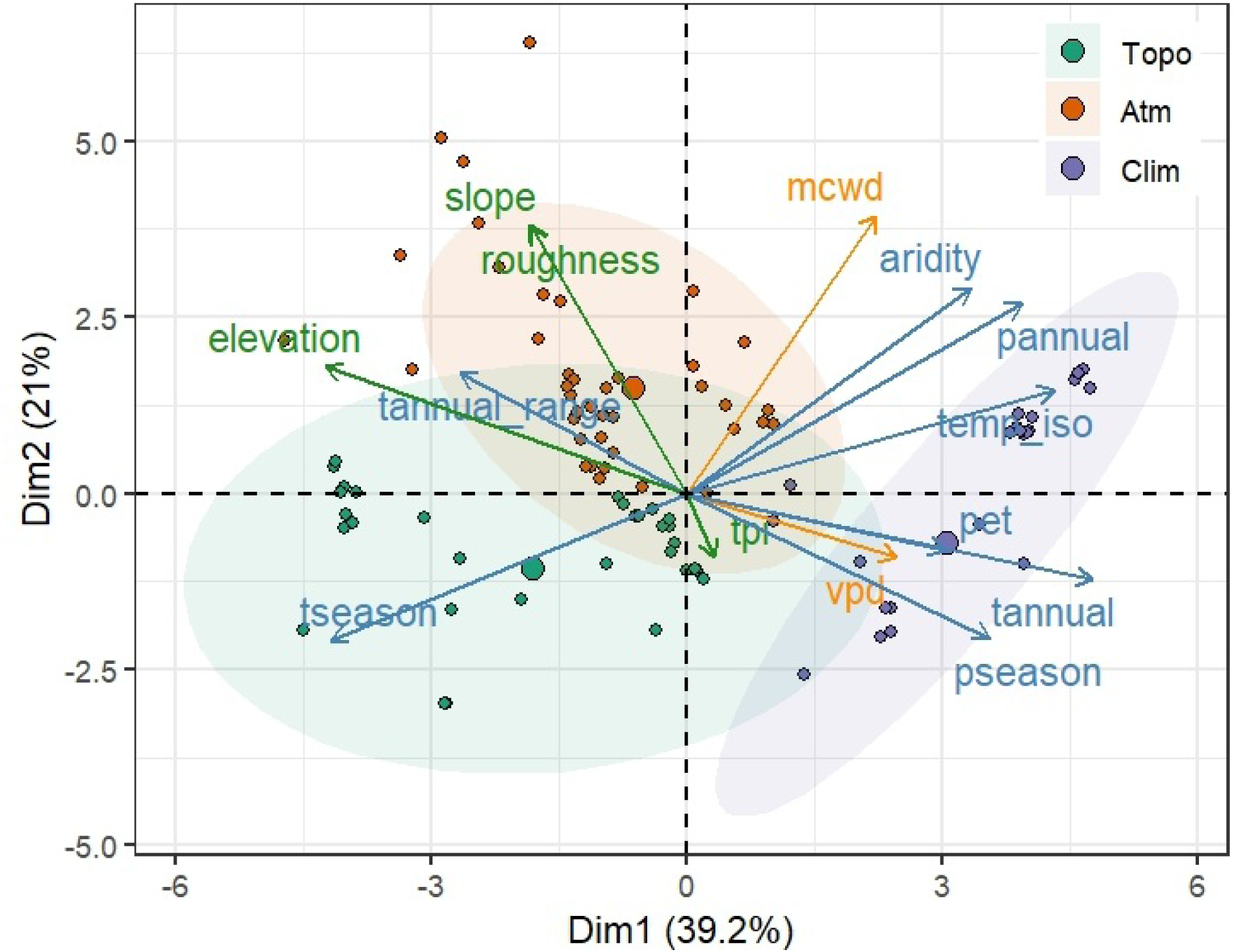
Environmental gradients structuring the AGB in the Caatinga (PCA–biplot). Biplot of principal components analysis with plots (dots) and environmental variables (arrows). Axis 1 (Dim1, 39.2%) organizes a hydroclimatic gradient dominated by annual precipitation, MCWD (accumulated water deficit; more negative values = deeper drought), VPD, and PET; axis 2 (Dim2, 21.0%) separates a topographic gradient (slope, rugosity, elevation, TPI). Arrow colors indicate thematic groups (orange = atmospheric, blue = climatic, green = topographic), and their length is proportional to the variable’s contribution to the axes. Ellipses represent envelopes (95%) of the plot groups in the PC space. Point size is proportional to AGB (log scale), demonstrating greater biomass in combinations of lower atmospheric demand/less drought and favorable topographic positions.

Our Random Forest model achieved robust predictive performance in estimating aboveground biomass (AGB) across the Caatinga. Under random *k*-fold cross-validation, the model explained 78% of the variance in observed biomass values (*R*^2^ = 0.78) with a root mean squared error (RMSE) of 15.4 Mg ha*^−^*^1^, indicating strong accuracy in reproducing local-scale variation. When evaluated through spatial cross-validation–which provides a more conservative measure of predictive skill by minimizing spatial autocorrelation–the model retained high explanatory power (*R*^2^ = 0.81) while showing a larger RMSE of 21 Mg ha*^−^*^1^, consistent with the increased difficulty of predicting across environmentally distinct regions. Residuals displayed no systematic bias or spatial structure, supporting the reliability of the predictions (Figures SI 1a,b). Collectively, these results demonstrate that the fitted Random Forest model captures both the magnitude and spatial heterogeneity of biomass across the biome, providing a solid foundation for spatial mapping and ecological interpretation of environmental controls.

Partial dependence plots reveal consistent nonlinear responses and thresholds (Figure 3). AGB sharply increases when MCWD is less negative than *<* 500 mm, rising from ∼20,000 Mg ha*^−^*^1^ under severe drought (*<* 700 mm) to *>* 35,000 Mg ha*^−^*^1^ in less water-limited conditions. Annual precipitation reinforces this pattern, with biomass gains up to ∼1,100–1,200 mm yr*^−^*^1^, beyond which AGB stabilizes. Conversely, atmospheric demand exerts a negative control: increases in PET (1,500–1,900 mm) or VPD (*>* 18,000 kPa) coincide with biomass declines of ∼5,000–7,000 Mg ha*^−^*^1^. The variable importance analysis further supports the functional responses revealed by partial dependence plots (Figures S2 a,b). Both metrics of importance (%IncMSE and IncNodePurity) consistently identify aridity, annual precipitation, topographic position (TPI), slope, ruggedness, MCWD, and VPD as the dominant predictors of AGB across the Caatinga.

**Figure 3:**
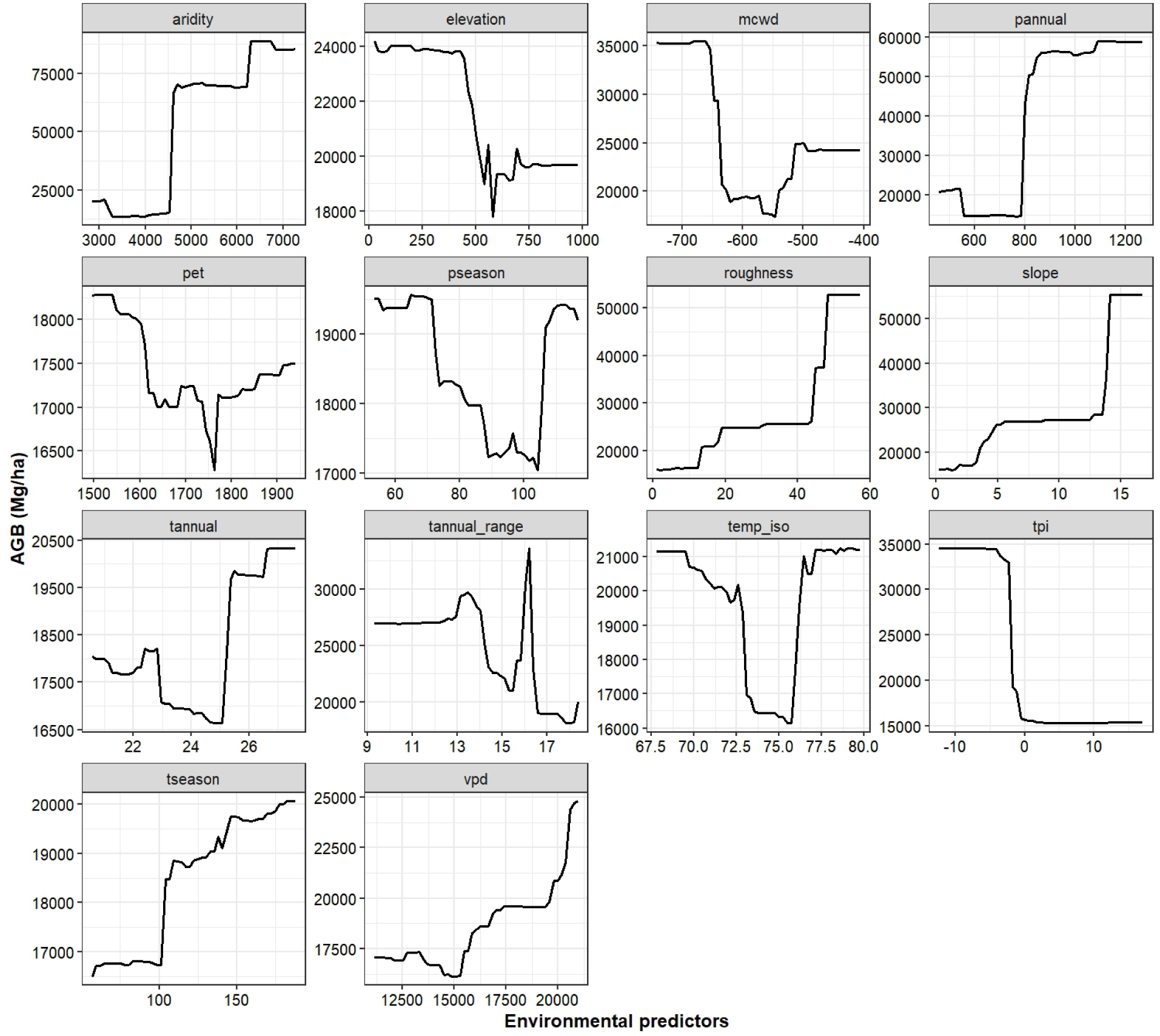
Partial responses of biomass to environmental predictors (Random Forests). Partial dependence curves show the marginal effect of each predictor on aboveground biomass (AGB, Mg ha*^−^*^1^), leaving the others within the observed distribution. Water and atmospheric factors exhibit clear thresholds: AGB increases sharply as MCWD becomes less negative (less severe drought) and annual precipitation increases to a saturation level, while higher VPD and PET reduce AGB. Topographic predictors additionally modulate stocks: roughness and slope tend to increase AGB (likely due to water retention/microclimate), while TPI indicates losses in convex/exposed positions. Thermal predictors (tannual, temp iso, tseason) exhibit nonlinear responses, with higher sensitivity bands. The curves reinforce that AGB is controlled by a regional hydrothermal axis and a relatively independent topographic axis, with mostly additive effects and ecologically plausible inflection points.

Aridity and precipitation appear as the strongest climatic drivers, underscoring the overarching role of water balance in structuring biomass distribution. Topographic variables (TPI, slope, ruggedness) rank immediately after, highlighting their capacity to locally buffer or amplify hydro-climatic stress. MCWD emerges as the leading drought metric, while VPD complements this by capturing atmospheric demand effects, consistent with the biomass declines observed at high evaporative loads. In contrast, variables such as isothermality, temperature seasonality, and potential evapotranspiration (PET) show comparatively weaker influence, acting more as modulators than as primary controls. These results corroborate the nonlinear thresholds described in the partial dependence plots: biomass accumulation accelerates when MCWD values are less negative, precipitation exceeds 1,100 mm, or ruggedness and slope reach critical thresholds, while atmospheric demand (high VPD, PET) and convex topographic positions constrain carbon storage. Together, the importance ranking and functional responses converge to depict a dual control system in which regional water availability and local topography jointly govern the resilience and spatial heterogeneity of biomass stocks in the Caatinga.

Topography further modulates these trends. Biomass rises steeply in rough terrains (*>*40 units of roughness), doubling relative to smoother surfaces, suggesting that micro-heterogeneity promotes soil accumulation and water retention. Slopes above ∼10° also support higher AGB, whereas convex landforms (positive TPI *>* 5) display sharp declines, with biomass falling from ∼30,000 Mg ha*^−^*^1^ to *<*15,000 Mg ha*^−^*^1^, consistent with exposure to higher evaporative stress. Elevation effects are segmented: biomass peaks at low (*<*200 m) and high (*>*600 m) sites but dips at intermediate altitudes, likely reflecting geomorphological transitions (Figure 3). Together, these results show that AGB in the Caatinga is structured by a dual control system: (i) a hydro-thermal axis where alleviation of seasonal drought (higher rainfall, less negative MCWD, lower PET/VPD) supports biomass accumulation, and (ii) an independent topographic axis where relief complexity redistributes resources and creates localized refugia. The absence of strong collinearity between axes highlights additive effects, with potential interactions only under extreme conditions––such as severe drought coinciding with exposed convex landforms.

### 3.2 Spatial distribution of AGB and environmental extremes

The spatial model reveals marked regional contrasts in aboveground biomass (AGB) across the Caatinga (Figure 4). Predicted stocks range from near-zero to *>* 120 Mg ha*^−^*^1^, with the lowest values (*<* 40 Mg ha*^−^*^1^) concentrated in central and interior depressions (≈8–12°S; 40–42°W). These zones coincide with highly negative MCWD (≤ −600 mm), elevated vapor pressure deficit (*>* 18,000 Pa), and pronounced aridity (Figure S3), conditions that sustain sparse vegetation and minimal carbon stocks. Intermediate values (40–80 Mg ha*^−^*^1^) dominate large portions of the northern (4–6°S; 37–39°W) and southern (13–15°S; 41–44°W) sectors, where precipitation exceeds 800–1,000 mm yr*^−^*^1^ and topographic heterogeneity (ruggedness *>*15, slopes *>*10°) enhances soil moisture retention (Figure S3). Localized maxima (*>* 100 Mg ha*^−^*^1^) emerge along eastern and southeastern boundaries (10–14°S; 36–38°W), associated with higher altitudes (*>* 500 m), steep slopes, and favorable hydroclimatic balances (less negative MCWD and moderate VPD). These areas act as environmental refugia where topography and climate jointly buffer seasonal drought, allowing forest patches with disproportionately high carbon density (Figure S3). Model uncertainty remains relatively low (≤ 6 Mg ha*^−^*^1^) over most of the domain, increasing only in transitional zones and highly heterogeneous terrains.

**Figure 4:**
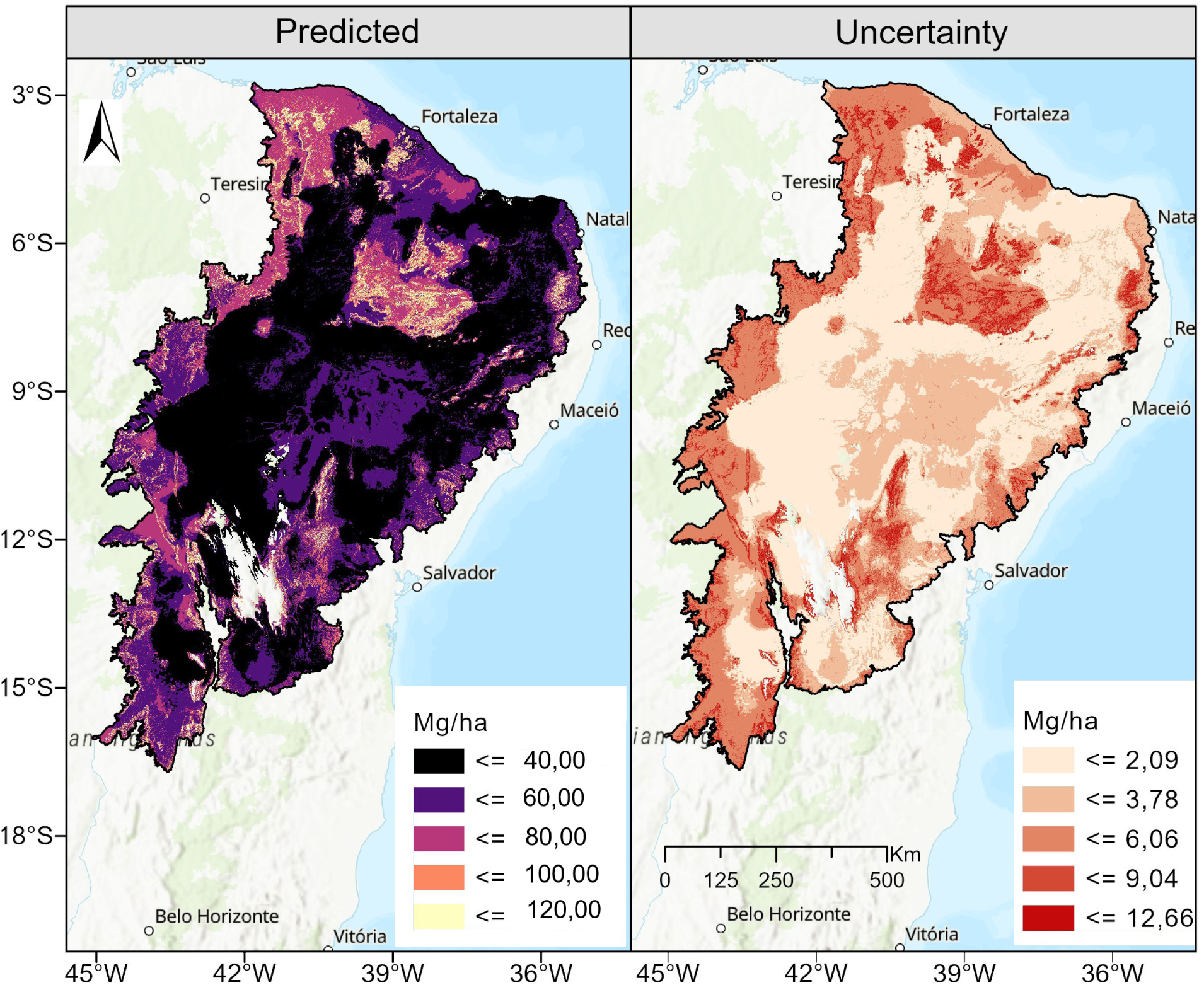
Spatial patterns of aboveground biomass (AGB) and model uncertainty in the Caatinga. Predicted AGB (left) ranges from *<* 40 Mg ha*^−^*^1^ in the central arid belt to localized hotspots exceeding 100 Mg ha*^−^*^1^ along eastern and southeastern gradients, where higher rainfall and topographic complexity create favorable conditions. Model uncertainty (right) remains low across most of the biome (*<* 6 Mg ha*^−^*^1^), with higher values confined to transitional zones and heterogeneous relief.

Patterns of environmental extremes confirm partially independent controls (Figure 5). The UpSet analysis shows that most extremes occur in isolation: severe MCWD (≤ −618 mm) in 39 plots and elevated ruggedness (≥ 15) in 37 plots dominate as the most recurrent single stressors. Other independent occurrences include high TPI (31 plots) and elevated aridity (26 plots). Pairwise intersections, such as MCWD with high rainfall or with VPD, appear in fewer than seven plots each, while three-way combinations are rare, restricted to ≤ 3 cases. This configuration suggests that environmental constraints on AGB operate largely additively, with only a handful of sites experiencing multiple simultaneous stressors.

**Figure 5:**
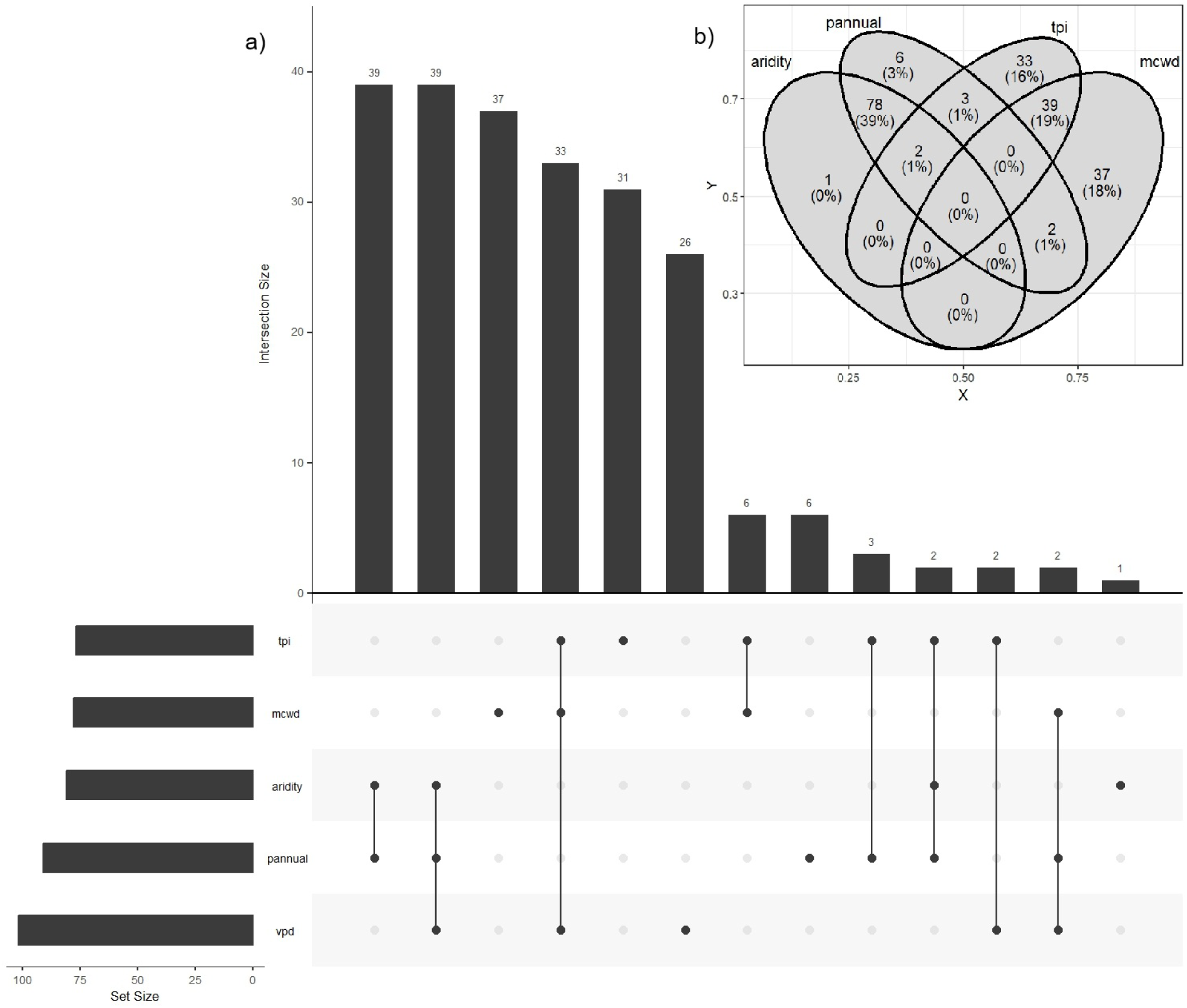
Co-occurrence of extreme environmental factors in Caatinga biomass plots. (**a**) UpSet diagram showing the intersections between plots that reach extreme environmental conditions. The largest bars correspond to isolated extremes (mainly severe MCWD and high roughness), while multiple co-occurrences are rare. (**b**) Venn diagram for four factors (aridity, annual precipitation, roughness, and MCWD), showing the relative frequencies of plots in each combination. The “MCWD-only” sector dominates (∼23% of plots), followed by roughness (∼11%). Multiple intersections (pairs or triplets) are in the minority, and there is no quadruple co-occurrence. Variables: aridity = aridity index; pannual = annual precipitation; roughness = terrain roughness; mcwd = Maximum Cumulative Water Deficit (more negative = more severe drought); vpd = vapor pressure deficit. Notes (thresholds used for binarization): Cuts (quartiles): Dryness ≥ 4500 — PAnnual. ≥ 790 —Roughness ≥ 14.75 — MCWD ≤ −618.6.

The Venn diagram reinforces this pattern: MCWD alone accounts for 69 plots (∼23% of the total), confirming its role as the dominant drought-related stressor (Figure 5). Ruggedness extremes occur in 33 plots (∼11%), underscoring topography as an independent local driver. High annual precipitation appears in only six plots (2%), while aridity extremes are virtually absent as isolated factors, usually co-occurring with MCWD. Triple or quadruple overlaps are negligible, indicating that combined severe drought, topographic exposure, and atmospheric demand rarely coincide in the Caatinga. Ecologically, this implies that carbon storage responds primarily to the depth of seasonal drought and secondarily to topographic regulation of water availability, while precipitation alone plays a minor role once MCWD and ruggedness are accounted for.

## 4 Discussion and conclusion

The strong influence of maximum cumulative water deficit (MCWD) on aboveground biomass (AGB) regulation reinforces the idea that, in seasonally dry tropical ecosystems such as the Caatinga, the severity of the dry season often acts as a more powerful ecological filter than total annual precipitation. In tropical forests and dryland ecosystems worldwide, prolonged droughts have already been identified as drivers of large tree mortality and progressive reductions in carbon stocks (e.g., de Meira Junior et al., 2020). When MCWD exceeds critical thresholds–such as the transition observed in our models between approximately −700 and −500 mm–plants experience extreme water stress, which suppresses photosynthetic rates, increases the risk of cavitation, and promotes leaf shedding or mortality. This turning point suggests that the Caatinga may be operating close to ecological tipping points, where relatively small changes in water balance can generate disproportionate impacts on biomass stocks.

In addition, atmospheric demand variables such as vapor pressure deficit (VPD) and potential evapotranspiration (PET) emerge as antagonistic regulators to water availability. Even under moderate precipitation, biomass accumulation may not occur if the atmosphere “pulls” too strongly on available water. Recent studies demonstrate that increases in VPD intensify soil desiccation and plant transpiration, while imposing stomatal constraints that limit carbon uptake (Novick et al., 2024; Stevens et al., 2021). In ecosystems already close to water deficit thresholds, this added atmospheric demand can shift systems into more critical regimes, underscoring how the VPD gradient observed in our models complements the drought control represented by MCWD.

Topography (ruggedness, slope, topographic position index, and elevation) emerges as a critical local modulator of biomass stocks, providing an important counterbalance: while macroclimatic gradients impose broad-scale limitations, terrain heterogeneity allows for the creation of microrefugia that buffer regional water stress. This role of topography is rarely captured in large-scale biomass models but has been emphasized in dryland ecosystems, where valleys, slopes, or areas of convergent flow favor water accumulation under stressful conditions (Siyum, 2020). In the Caatinga, such micro-structures may represent refugia where more demanding species persist and accumulate higher biomass.

These findings have critical implications for carbon stock estimation and for projections under altered climate regimes. Regional and global models that ignore local topographic effects or the nonlinear drought thresholds may underestimate the vulnerability of the Caatinga to extreme events (J. L. S. E. Silva et al., 2019). Although substantial carbon stocks have been documented in the biome (Althoff et al., 2018; Pereira Júnior et al., 2016), high-resolution spatial studies that jointly incorporate topography and atmospheric demand remain scarce. Under future scenarios of intensified drought and increased evaporative demand–as projected by many climate models–the tolerance margin of Caatinga forests may be surpassed, leading to abrupt biomass declines and carbon release to the atmosphere (Cavalcante et al., 2023). The predominance of single environmental extremes acting independently across plots (rather than multiple stressors simultaneously) further suggests that biomass regulation is largely additive, with limited synergistic interactions–except under extreme cases. This highlights that biomass control in the Caatinga is not centralized in a single driver but results from a web of ecological forces operating across different scales (Souza et al., 2021). Recognizing this multifactorial structure is essential for restoration planning, forest management, and climate modeling: areas with favorable topography should be prioritized as “hotspots” of resistance to climate stress, while zones under high evaporative demand may require targeted interventions for conservation.

Our analysis demonstrates that the Caatinga, although often perceived as a low-biomass ecosystem, harbors carbon-dense refugia where topography and water regulation interact synergistically. This spatial heterogeneity underscores that biomass is not simply or uniformly distributed but responds to a combination of climatic gradients, atmospheric demand, and local microenvironments (Castanho et al., 2020; A. C. Silva & Souza, 2022). In seasonal tropical forests more broadly, carbon stocks vary widely: for instance, Maia et al., 2020 showed that Brazilian tropical seasonal forests act as significant carbon sinks even under high climatic variability. In less extreme ecosystems, the importance of drought tolerance is well recognized (Haghpanah et al., 2024; Shao et al., 2024); yet in the Caatinga, our evidence of abrupt thresholds (such as the transition from MCWD ≈ −700 to −500 mm) suggests that this biome operates near viability limits in many locations.

Comparisons with dry forests elsewhere reinforce the mechanisms identified here. Across tropical forests exposed to prolonged drought, increases in both water deficit and VPD have been linked to rising tree mortality and declining biomass stocks (e.g., Aguirre-Gutiérrez et al., 2020; Bauman et al., 2022). A pan-tropical study (Bauman et al., 2022) demonstrated that mortality risk increases as VPD rises, effectively shortening carbon residence time in trees (i.e., accelerating biomass losses) and compromising sink stability. This evidence indicates that while MCWD represents the primary hydraulic bottleneck in our model, VPD and atmospheric demand should not be underestimated– particularly under global warming scenarios where VPD is expected to intensify.

The role of topography as a local modulator functions as a buffer, enabling some locations to escape the broader limits imposed by regional aridity. Such microrefugial dynamics are well documented in ecological studies (e.g., Lenoir et al., 2016; Denney et al., 2020), though rarely mapped at large scales: complex terrain can generate areas of soil and water accumulation, subtle microclimatic differences (cooler temperatures, reduced wind exposure), and lower stress for seedlings. These sites may therefore sustain communities with more demanding woody species. This insight highlights the need for future biomass and carbon models to include explicit topographic metrics (ruggedness, slope, and TPI), rather than relying solely on coarse terrain proxies.

From a practical and restoration perspective, our results yield clear implications. First, areas with favorable terrain (valleys, sheltered slopes) should be prioritized as targets for restoration or conservation, as they are more resilient to water stress. Conversely, flat or highly exposed areas, particularly those already subjected to severe water deficits, warrant special management attention (e.g., soil conservation measures, supplementary irrigation, or drought-tolerant species selection). Moreover, regional carbon stock estimates that overlook local heterogeneity and nonlinear drought thresholds are likely to overestimate ecosystem stability under climate change scenarios (C. D. Allen et al., 2015; Li et al., 2025). Looking ahead, if more severe droughts and higher VPD levels become widespread–as projected in many climate models–large portions of the Caatinga’s biomass stocks may be at risk.

Taken together, our results reveal that aboveground biomass in the Caatinga is governed by a dual regime of control: (i) a hydroclimatic axis, where water availability and drought severity impose nonlinear thresholds (AGB rises sharply when MCWD becomes less negative than −500 mm and annual precipitation surpasses ∼1,100 mm, while high VPD and PET reduce stocks); and (ii) an independent topographic axis, where slope, ruggedness, and landform position create microrefugia that buffer evaporative stress and sustain high carbon density. The spatial model performed robustly (*R*^2^ = 0.81 under spatial cross-validation; RMSE = 21 Mg ha*^−^*^1^), enabling the mapping of a mosaic of low-biomass nuclei interspersed with high-biomass refugia in complex terrains. This perspective rejects the notion of a homogeneous biome and reframes the Caatinga as a heterogeneous, dynamic carbon reservoir, particularly sensitive to intensifying droughts and increasing atmospheric demand under global warming. By providing both a predictive map and a mechanistic framework for the combined roles of climate, atmospheric stressors, and topography, our study offers operational foundations for prioritizing conservation and restoration areas, calibrating inventories, and informing public policy for mitigation and adaptation. Future advances should integrate controlled experiments, explicit modeling of interactions (MCWD × VPD× topography), and targeted inventories in high-biomass enclaves to reduce uncertainties and guide landscape-scale decision-making.

## Authors’ contributions

Brhenda S. Lozado, Robson B. de Lima, Cinthia P. de Oliveira, Alessandro de Paula, Patrícia A. B. Barreto-Garcia, Odair L. Lemos, Ana Luisa L. Pereira conceptualised the study. Brhenda S. Lozado, Robson B. de Lima, Cinthia P. de Oliveira, Alessandro de Paula, Patrícia A. B. Barreto-Garcia, Odair L. Lemos, Ana Luisa L. Pereira, Frans Pareyn, Peter W. Moonlight, Domingos Cardoso, Elmar Veenendaal, Luciano P. de Queiroz, Priscyla M. S. Rodrigues, Rubens M. dos Santos, Tiina Sarkinen, Toby Pennington, Oliver L. Phillips collected the data. Brhenda S. Lozado, Robson B. de Lima, Cinthia P. de Oliveira, Emanuel A. Silva analysed the data and produced the figures. Brhenda S. Lozado, Robson B. de Lima and Cinthia P. de Oliveira wrote the initial manuscript. Rinaldo L. C. Ferreira and José A. Aleixo da Silva contributed to subsequent iterations of the manuscript. Allauthors approved the final version for submission.

## Acknowledgements

We thank the ForestPlots.net, Inema and Caatinga forest management network teams for permission and availability of data to conduct the research. This project was supported by ForestPlots.net approved Research Project 142: “Mapping the biomass and carbon stock, volume and diversity in tropical forests”, and the Programa de Pós-graduação em Ciências Florestais da Universidade Estadual do Sudoeste da Bahia (PPGCIFLOR/UESB).

## Fundingin Formation

UK NERC Newton Fund project “Nordeste” (NE/N012550/1). The authors thank the Fundação de Amparo da Pesquisa da Bahia (FAPESB; Processes nQ 084.0508.2023.0004282-06) and Conselho Nacional de Desenvolvimento Científico e Tecnológico (Processes 301432/2022-8) for financial support in a research grant for the first and last author of this work.

## Supporting information

**Figure 1.**
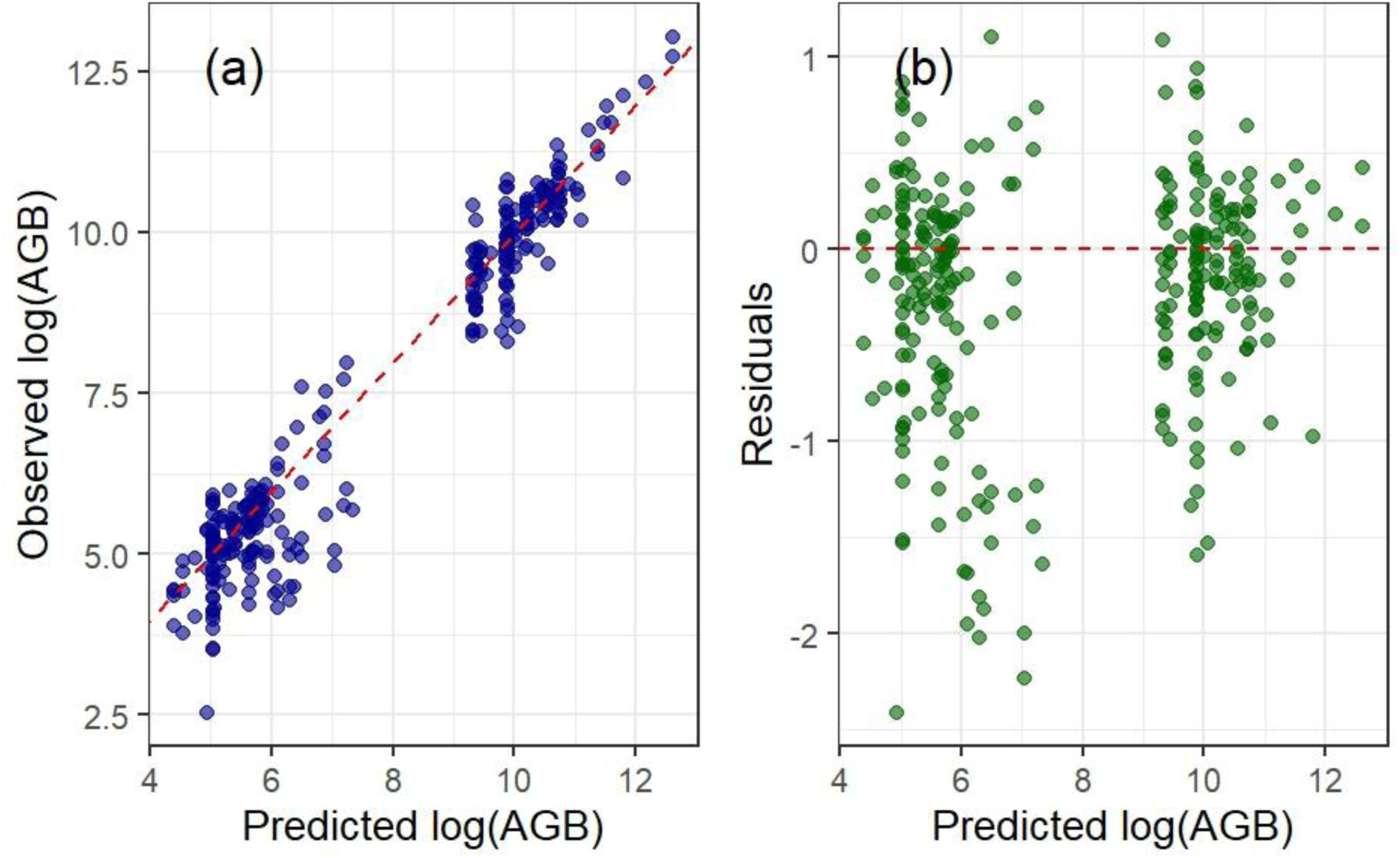
Model performance of Random Forest predictions of aboveground biomass (AGB) in the Caatinga. (a) Relationship between observed and predicted AGB values (log-transformed) after excluding three extreme outliers. The dashed red line represents the 1:1 relationship. (b) Residuals plotted against predicted AGB (log-transformed), with the dashed red line indicating zero residuals. The model achieved strong predictive performance (R² = 0.78 under random cross-validation and R² = 0.81 under spatial cross-validation), with RMSE values of 15.4 Mg ha⁻¹and 21 Mg ha⁻¹, respectively, confirming the reliability of the Random Forest framework for capturing biomass variation across the biome.

**Figure 2.**
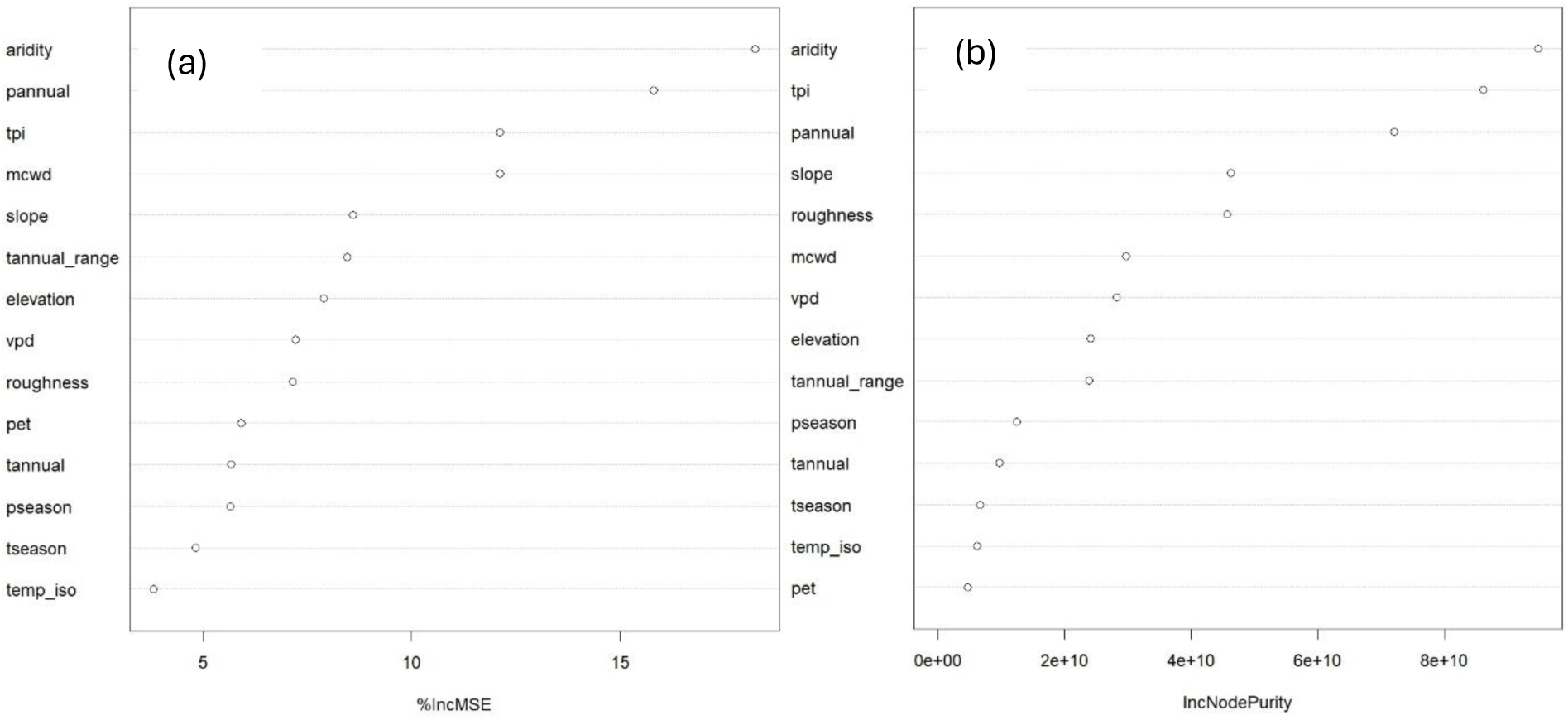
Variable importance of environmental predictors in Random Forest models of aboveground biomass (AGB) in the Caatinga. (a) Predictor importance measured by the percentage increase in mean squared error (%IncMSE), which reflects the reduction in model accuracy when each variable is permuted. Higher values indicate stronger contributions to predictive performance. (b) Predictor importance measured by the increase in node purity (IncNodePurity), representing the total reduction in residual variance explained by each split across all trees. Both metrics consistently highlight **aridity index, precipitation, topographic position index (TPI), maximum cumulative water deficit (MCWD), and slope** as dominant predictors of AGB variation, while variables such as temperature isothermality (*temp_iso*) and temperature seasonality (*tseason*) contributed minimally. Together, these results emphasize the dual control of hydroclimatic stressors and terrain heterogeneity in shaping biomass distribution across the Caatinga.

**Figure 3.**
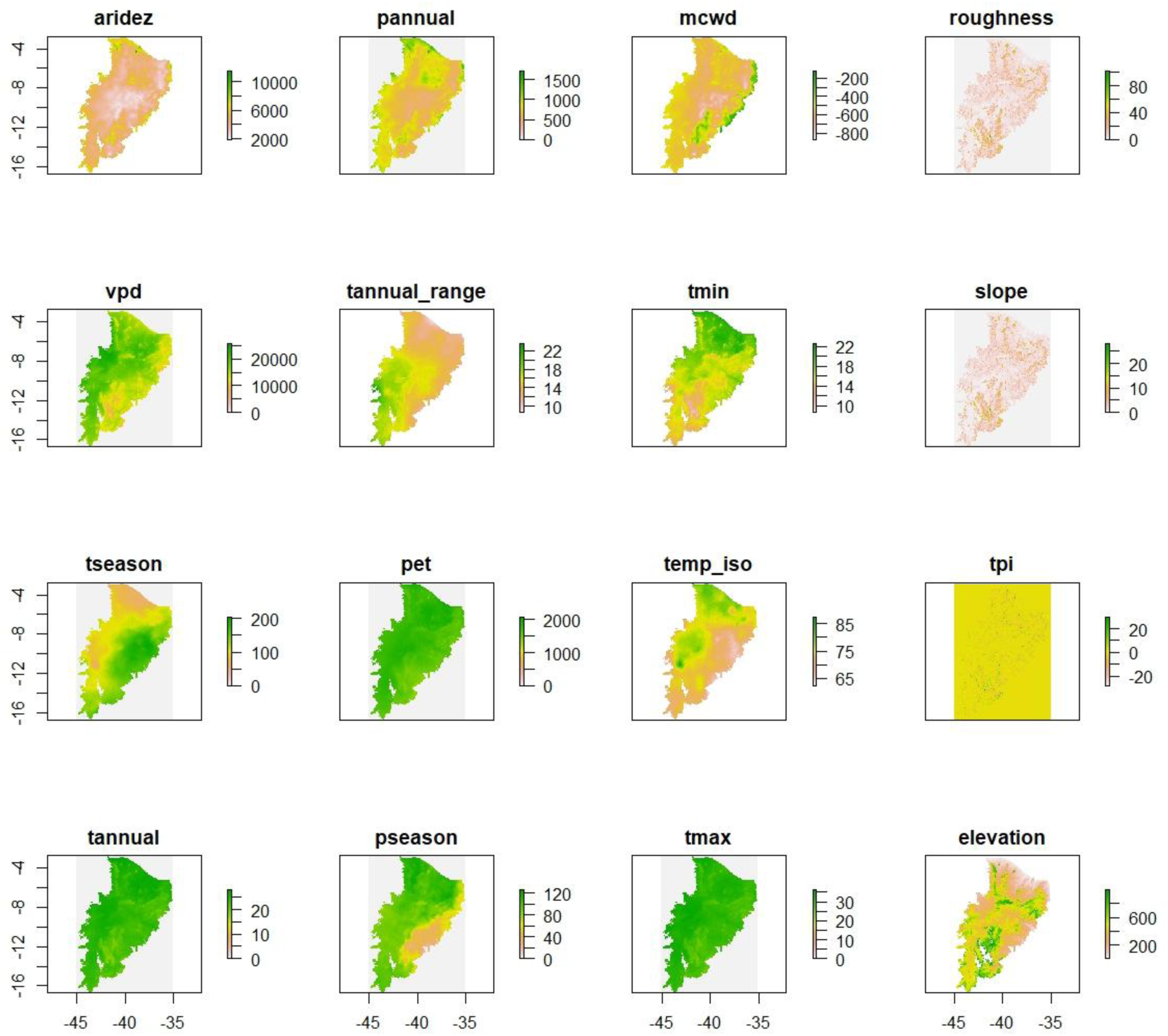
Spatial distribution of environmental predictors used in the modeling of aboveground biomass (AGB) in the Caatinga. Maps illustrate the spatial patterns of climatic, atmospheric, and topographic predictors at 30 arc-seconds (∼1 km) resolution. Climatic variables include mean annual precipitation (*pannual*), mean annual temperature (*tannual*), annual temperature range (*tannual_range*), temperature seasonality (*tseason*), precipitation seasonality (*pseason*), maximum temperature (*tmax*), and minimum temperature (*tmin*). Atmospheric stressors comprise vapor pressure deficit (*vpd*), potential evapotranspiration (*pet*), aridity index (*aridez*), and maximum cumulative water deficit (*mcwd*). Topographic variables include elevation (*elevation*), slope (*slope*), roughness (*roughness*), and topographic position index (*tpi*). Together, these predictors capture large-scale hydroclimatic gradients and local terrain heterogeneity, which jointly modulate water availability, atmospheric demand, and microclimatic conditions influencing biomass distribution across the biome.

## References

1. Abatzoglou, J. T., Dobrowski, S. Z., Parks, S. A., & Hegewisch, K. C. (2018). TerraClimate, a highresolution global dataset of monthly climate and climatic water balance from 1958–2015. Scientific Data, 5(1), 170191. 10.1038/sdata.2017.191

2. Aguirre-Gutiérrez, J., Malhi, Y., Lewis, S. L., Fauset, S., Adu-Bredu, S., Affum-Baffoe, K., Baker, T. R., Gvozdevaite, A., Hubau, W., Moore, S., Peprah, T., Ziemińska, K., Phillips, O. L., & Oliveras, I. (2020). Long-term droughts may drive drier tropical forests towards increased functional, taxonomic and phylogenetic homogeneity. Nature Communications, 11(1), 3346. 10.1038/s41467-020-16973-4

3. Allen, C. D., Breshears, D. D., & McDowell, N. G. (2015). On underestimation of global vulnerability to tree mortality and forest die-off from hotter drought in the Anthropocene. Ecosphere, 6(8), art129. 10.1890/ES15-00203.1

4. Allen, K., Dupuy, J. M., Gei, M. G., Hulshof, C., Medvigy, D., Pizano, C., Salgado-Negret, B., Smith, C. M., Trierweiler, A., Bloem, S. J. V., Waring, B. G., Xu, X., & Powers, J. S. (2017). Will seasonally dry tropical forests be sensitive or resistant to future changes in rainfall regimes? Environmental Research Letters, 12(2), 023001. 10.1088/1748-9326/aa5968

5. Althoff, T. D., Menezes, R. S. C., Pinto, A. de S., Pareyn, F. G. C., Carvalho, A. L. de, Martins, J. C. R., de Carvalho, E. X., Silva, A. S. A. da, Dutra, E. D., & Sampaio, E. V. de S. B. (2018). Adaptation of the century model to simulate C and N dynamics of Caatinga dry forest before and after deforestation. Agriculture, Ecosystems & Environment, 254, 26–34. 10.1016/j.agee.2017.11.016

6. Alves Rodrigues Pinheiro, E., Metselaar, K., de Jong van Lier, Q., & de Araújo, J. C. (2016). Importance of soil-water to the Caatinga biome, Brazil. Ecohydrology, 9(7), 1313–1327. 10.1002/eco.1728

7. Amatulli, G., Domisch, S., Tuanmu, M.-N., Parmentier, B., Ranipeta, A., Malczyk, J., & Jetz, W. (2018). A suite of global, cross-scale topographic variables for environmental and biodiversity modeling. Scientific Data, 5(1), 180040. 10.1038/sdata.2018.40

8. Amorim, I. L. de, Sampaio, E. V. S. B., & Araújo, E. de L. (2005). Flora e estrutura da vegetação arbustivo-arbórea de uma área de caatinga do Seridó, RN, Brasil. Acta Botanica Brasilica, 19(3), 615–623. 10.1590/S0102-33062005000300023

9. Anderson, K. E., Glenn, N. F., Spaete, L. P., Shinneman, D. J., Pilliod, D. S., Arkle, R. S., McIlroy, S. K., & Derryberry, D. R. (2018). Estimating vegetation biomass and cover across large plots in shrub and grass dominated drylands using terrestrial lidar and machine learning. Ecological Indicators, 84, 793–802. 10.1016/j.ecolind.2017.09.034

10. Apgaua, D. M. G., Tng, D. Y. P., & Laurance, S. G. W. (2022). Tropical wet and dry forest tree species exhibit contrasting hydraulic architecture. Flora, 291, 152072. 10.1016/j.flora.2022.152072

11. Araujo, H. F. P., Canassa, N. F., Machado, C. C. C., & Tabarelli, M. (2023). Human disturbance is the major driver of vegetation changes in the Caatinga dry forest region. Scientific Reports, 13(1), 18440. 10.1038/s41598-023-45571-9

12. Baccini, A., & Asner, G. P. (2013). Improving pantropical forest carbon maps with airborne LiDAR sampling. Carbon Management, 4(6), 591–600. 10.4155/cmt.13.66

13. Baccini, A., Walker, W., Carvalho, L., Farina, M., Sulla-Menashe, D., & Houghton, R. A. (2017). Tropical forests are a net carbon source based on aboveground measurements of gain and loss. Science, 358(6360), 230–234. 10.1126/science.aam5962

14. Barkhordarian, A., Saatchi, S. S., Behrangi, A., Loikith, P. C., & Mechoso, C. R. (2019). A Recent Systematic Increase in Vapor Pressure Deficit over Tropical South America. Scientific Reports, 9(1). 10.1038/s41598-019-51857-8

15. Bauman, D., Fortunel, C., Delhaye, G., Malhi, Y., Cernusak, L., Bentley, L., Rifai, S., Aguirre-Gutiérrez, J., Oliveras, I., Phillips, O., McNellis, B., Bradford, M., Laurance, S., Hutchinson, M. F., Dempsey, R., Santos-Andrade, P., Ninantay-Rivera, H., Paucar, J., & McMahon, S. (2022). Tropical tree mortality has increased with rising atmospheric water stress. Nature, 1–6. 10.1038/s41586-022-04737-7

16. Biomas — IBGE. (n.d.). Retrieved June 17, 2024. ibge.gov.br/…/15842-biomas.html

17. Calhoun, P., Hallett, M., Su, X., Cafri, G., Levine, R., & Fan, J. (2020). Random forest with acceptance–rejection trees. Computational Statistics, 35. 10.1007/s00180-019-00929-4

18. Campanharo, W., & Silva-Junior, C. (2019). Maximum Cumulative Water Deficit – MCWD: A R language script. 10.5281/zenodo.2652629

19. Campos, S., Mendes, K. R., da Silva, L. L., Mutti, P. R., Medeiros, S. S., Amorim, L. B., dos Santos, C. A. C., Perez-Marin, A. M., Ramos, T. M., Marques, T. V., Lucio, P. S., Costa, G. B., Santos e Silva, C. M., & Bezerra, B. G. (2019). Closure and partitioning of the energy balance in a preserved area of a Brazilian seasonally dry tropical forest. Agricultural and Forest Meteorology, 271, 398–412. 10.1016/j.agrformet.2019.03.018

20. Castanho, A. D. A., Coe, M., Andrade, E. M., Walker, W., Baccini, A., Campos, D. A., & Farina, M. (2020). A close look at above ground biomass of a large and heterogeneous Seasonally Dry Tropical Forest—Caatinga in North East of Brazil. Anais Da Academia Brasileira de Ciências, 92, e20190282. 10.1590/0001-3765202020190282

21. Cavalcante, A. M. B., Sampaio, A. C. P., Duarte, A. S., & Santos, M. A. F. D. (2023). Impacts of climate change on the potential distribution of epiphytic cacti in the Caatinga biome, Brazil. Anais Da Academia Brasileira de Ciências, 95, e20200904. 10.1590/0001-3765202320200904

22. Chave, J., Réjou-Méchain, M., Búrquez, A., Chidumayo, E., Colgan, M. S., Delitti, W. B. C., Duque, A., Eid, T., Fearnside, P. M., Goodman, R. C., Henry, M., Martínez-Yrízar, A., Mugasha, W. A., Muller-Landau, H. C., Mencuccini, M., Nelson, B. W., Ngomanda, A., Nogueira, E. M., Ortiz-Malavassi, E., … Vieilledent, G. (2014). Improved allometric models to estimate the aboveground biomass of tropical trees. Global Change Biology, 20(10), 3177–3190. 10.1111/gcb.12629

23. Costa, A. C., Rowland, L., Oliveira, R., Oliveira, A., Binks, O., Salmon, Y., Vasconcelos, S., Silva Junior, J., Ferreira, L., Poyatos, R., Mencuccini, M., & Meir, P. (2017). Stand dynamics modulate water cycling and mortality risk in droughted tropical forest. Global Change Biology, 24. 10.1111/gcb.13851

24. Cutler, F. original by L. B. and A., & Wiener, R. port by A. L. & M. (2022). randomForest: Breiman and Cutler’s Random Forests for Classification and Regression (Version 4.7-1.1) [Computer software]. CRAN.R-project.org/package=randomForest

25. Da Silva, J. M., Leal, I., & Tabarelli, M. (2017). Caatinga: The Largest Tropical Dry Forest Region in South America. In Caatinga: The Largest Tropical Dry Forest Region in South America. 10.1007/978-3-319-68339-3

26. de Meira Junior, M. S., Pinto, J. R. R., Ramos, N. O., Miguel, E. P., Gaspar, R. de O., & Phillips, O. L. (2020). The impact of long dry periods on the aboveground biomass in a tropical forest: 20 years of monitoring. Carbon Balance and Management, 15(1), 12. 10.1186/s13021-020-00147-2

27. Denney, D. A., Jameel, M. I., Bemmels, J. B., Rochford, M. E., & Anderson, J. T. (2020). Small spaces, big impacts: Contributions of micro-environmental variation to population persistence under climate change. AoB PLANTS, 12(2), plaa005. 10.1093/aobpla/plaa005

28. DRYFLOR, Banda-R, K., Delgado-Salinas, A., Dexter, K. G., Linares-Palomino, R., Oliveira-Filho, A., Prado, D., Pullan, M., Quintana, C., Riina, R., Rodríguez M., G. M., Weintritt, J., Acevedo-Rodríguez, P., Adarve, J., Álvarez, E., Aranguren B., A., Arteaga, J. C., Aymard, G., Castaño, A., … Pennington, R. T. (2016). Plant diversity patterns in neotropical dry forests and their conservation implications. Science, 353(6306), 1383–1387. 10.1126/science.aaf5080

29. Ferraz, A., Saatchi, S., Xu, L., Hagen, S., Chave, J., Yu, Y., Meyer, V., Garcia, M., Silva, C., Roswintiart, O., Samboko, A., Sist, P., Walker, S., Pearson, T. R. H., Wijaya, A., Sullivan, F. B., Rutishauser, E., Hoekman, D., & Ganguly, S. (2018). Carbon storage potential in degraded forests of Kalimantan, Indonesia. Environmental Research Letters, 13(9), 095001. 10.1088/1748-9326/aad782

30. ForestPlots.net, Blundo, C., Carilla, J., Grau, R., Malizia, A., Malizia, L., Osinaga-Acosta, O., Bird, M., Bradford, M., Catchpole, D., Ford, A., Graham, A., Hilbert, D., Kemp, J., Laurance, S., Laurance, W., Ishida, F. Y., Marshall, A., Waite, C., … Tran, H. D. (2021). Taking the pulse of Earth’s tropical forests using networks of highly distributed plots. Biological Conservation, 260, 108849. 10.1016/j.biocon.2020.108849

31. Gauthey, A., & Gardner, A. (2024). On the importance of vapor pressure deficit for the determination of the photosynthetic temperature optimum in tropical trees. New Phytologist, 244(4), 1119–1121. 10.1111/nph.20041

32. Genuer, R., & Poggi, J.-M. (2020). Random Forests with R. Springer International Publishing. 10.1007/978-3-030-56485-8

33. Haghpanah, M., Hashemipetroudi, S., Arzani, A., & Araniti, F. (2024). Drought Tolerance in Plants: Physiological and Molecular Responses. Plants, 13(21), 2962. 10.3390/plants13212962

34. Hijmans, R. J., Etten, J. van, Sumner, M., Cheng, J., Baston, D., Bevan, A., Bivand, R., Busetto, L., Canty, M., Fasoli, B., Forrest, D., Ghosh, A., Golicher, D., Gray, J., Greenberg, J. A., Hiemstra, P., Hingee, K., Karney, C., … Wueest, R. (2021). raster: Geographic Data Analysis and Modeling (Version 3.4-13) [Computer software]. CRAN.R-project.org/package=raster

35. Husson, F., Josse, J., Le, S., & Mazet, J. (2025). FactoMineR: Multivariate Exploratory Data Analysis and Data Mining (Version 2.12) [Computer software]. cran.r-project.org/web/packages/FactoMineR/index

36. Karger, D. N., Conrad, O., Böhner, J., Kawohl, T., Kreft, H., Soria-Auza, R. W., Zimmermann, N. E., Linder, H. P., & Kessler, M. (2017). Climatologies at high resolution for the earth’s land surface areas. Scientific Data, 4(1), 170122. 10.1038/sdata.2017.122

37. Lenoir, J., Hattab, T., & Pierre, G. (2016). Climatic microrefugia under anthropogenic climate change: Implications for species redistribution. Ecography, 40. 10.1111/ecog.02788

38. Lewis, S. L., Sonké, B., Sunderland, T., Begne, S. K., Lopez-Gonzalez, G., van der Heijden, G. M. F., Phillips, O. L., Affum-Baffoe, K., Baker, T. R., Banin, L., Bastin, J.-F., Beeckman, H., Boeckx, P., Bogaert, J., De Cannière, C., Chezeaux, E., Clark, C. J., Collins, M., Djagbletey, G., … Zemagho, L. (2013). Above-ground biomass and structure of 260 African tropical forests. Philosophical Transactions of the Royal Society B: Biological Sciences, 368(1625), 20120295. 10.1098/rstb.2012.0295

39. Li, T., Wang, Q., Wang, H., Wang, J., Li, X., Liao, Z., Luo, P., Lai, C., Liu, Y., & Jian, Y. (2025). Increases in forest carbon stocks of up to 32% by 2100 across the subalpine forests of the Tibetan Plateau. Ecological Indicators, 176, 113675. 10.1016/j.ecolind.2025.113675

40. Liang, J., Gamarra, J.G.P., Picard, N. et al. (2022). Co-limitation towards lower latitudes shapes global forest diversity gradients. Nature Ecology & Evolution, 6, 1423–1437. 10.1038/s41559-022-01831-x

41. Maia, V., Santos, A., de Aguiar-Campos, N., Souza, C., Coutinho Freitas de Oliveira, M., Coelho, P., Morel, J., Costa, L., Farrapo, C., Fagundes, N., Pires, G., Santos, P., Gianasi, F., Silva, W., Oliveira, F., Girardelli, D., Araújo, F., Vilela, T., Pereira, R., & Santos, R. (2020). The carbon sink of tropical seasonal forests in southeastern Brazil can be under threat. Science Advances, 6, eabd4548. 10.1126/sciadv.abd4548

42. Mattos, C. R. C., Hirota, M., Oliveira, R. S., Flores, B. M., Miguez-Macho, G., Pokhrel, Y., & Fan, Y. (2023). Double stress of waterlogging and drought drives forest–savanna coexistence. Proceedings of the National Academy of Sciences, 120(33), e2301255120. 10.1073/pnas.2301255120

43. Mendes, K. R., Campos, S., da Silva, L. L., Mutti, P. R., Ferreira, R. R., Medeiros, S. S., Perez-Marin, A. M., Marques, T. V., Ramos, T. M., de Lima Vieira, M. M., Oliveira, C. P., Gonçalves, W. A., Costa, G. B., Antonino, A. C. D., Menezes, R. S. C., Bezerra, B. G., & Santos e Silva, C. M. (2020). Seasonal variation in net ecosystem CO_2_ exchange of a Brazilian seasonally dry tropical forest. Scientific Reports, 10(1), 9454. 10.1038/s41598-020-66415-w

44. Miles, L., Newton, A. C., DeFries, R. S., Ravilious, C., May, I., Blyth, S., Kapos, V., & Gordon, J. E. (2006). A global overview of the conservation status of tropical dry forests. Journal of Biogeography, 33(3), 491–505. 10.1111/j.1365-2699.2005.01424.x

45. Novick, K. A., Ficklin, D. L., Grossiord, C., Konings, A. G., Martínez-Vilalta, J., Sadok, W., Trugman, A. T., Williams, A. P., Wright, A. J., Abatzoglou, J. T., Dannenberg, M. P., Gentine, P., Guan, K., Johnston, M. R., Lowman, L. E. L., Moore, D. J. P., & McDowell, N. G. (2024). The impacts of rising vapour pressure deficit in natural and managed ecosystems. Plant, Cell & Environment, 47(9), 3561–3589. 10.1111/pce.14846

46. Oliveira, C., Luiz, R., Silva, J., Lima, R., Silva, E., da Silva, A., Lucena, J., Tavares dos Santos, N., Lopes, I., Pessoa, M., & Melo, C. (2021). Modeling and Spatialization of Biomass and Carbon Stock Using LiDAR Metrics in Tropical Dry Forest, Brazil. Forests, 12, 1–17. 10.3390/f12040473

47. Pereira Júnior, L. R., Andrade, E. M. de, Palácio, H. A. de Q., Raymer, P. C. L., Ribeiro Filho, J. C., & Pereira, F. J. S. (2016). Carbon stocks in a tropical dry forest in Brazil. Revista Cîencia Agronômica, 47, 32–40. 10.5935/1806-6690.20160004

48. Powers, J. S., Feng, X., Sanchez-Azofeifa, A., & Medvigy, D. (2018). Focus on tropical dry forest ecosystems and ecosystem services in the face of global change. Environmental Research Letters, 13(9), 090201. 10.1088/1748-9326/aadeec

49. Roberts, D. R., Bahn, V., Ciuti, S., Boyce, M. S., Elith, J., Guillera-Arroita, G., Hauenstein, S., Lahoz-Monfort, J. J., Schröder, B., Thuiller, W., Warton, D. I., Wintle, B. A., Hartig, F., & Dormann, C. F. (2017). Cross-validation strategies for data with temporal, spatial, hierarchical, or phylogenetic structure. Ecography, 40(8), 913–929. 10.1111/ecog.02881

50. Saatchi, S., Malhi, Y., Zutta, B., Buermann, W., Anderson, L. O., Araujo, A. M., Phillips, O. L., Peacock, J., ter Steege, H., Lopez Gonzalez, G., Baker, T., Arroyo, L., Almeida, S., Higuchi, N., Killeen, T., Monteagudo, A., Neill, D., Pitman, N., Prieto, A., … Ramírez, H. A. (2009). Mapping landscape scale variations of forest structure, biomass, and productivity in Amazonia [Preprint]. 10.5194/bgd-6-5461-2009

51. Saatchi, S. S., Harris, N. L., Brown, S., Lefsky, M., Mitchard, E. T. A., Salas, W., Zutta, B. R., Buermann, W., Lewis, S. L., Hagen, S., Petrova, S., White, L., Silman, M., & Morel, A. (2011). Benchmark map of forest carbon stocks in tropical regions across three continents. Proceedings of the National Academy of Sciences, 108(24), 9899–9904. 10.1073/pnas.1019576108

52. Sampaio, E., Gasson, P., Baracat, A., Cutler, D., Pareyn, F., & Lima, K. C. (2010). Tree biomass estimation in regenerating areas of tropical dry vegetation in northeast Brazil. Forest Ecology and Management, 259(6), 1135–1140. 10.1016/j.foreco.2009.12.028

53. Santos, H. G. dos (with Embrapa Solos). (2018). Sistema brasileiro de classificação de solos (5a edição revista e ampliada). Embrapa.

54. Schönbeck, L., Schuler, P., Lehmann, M., Mas, E., Mekarni, L., Pivovaroff, A., Turberg, P., & Grossiord, C. (2022). Increasing temperature and vapour pressure deficit lead to hydraulic damages in the absence of soil drought. Plant, Cell & Environment, 45, 3275–3289. 10.1111/pce.14425

55. Shao, X., Zhang, Y., Ma, N., Zhang, X., Tian, J., Xu, Z., & Liu, C. (2024). Drought-induced ecosystem resistance and recovery observed at 118 flux tower stations across the globe. Agricultural and Forest Meteorology, 356, 110170. 10.1016/j.agrformet.2024.110170

56. Silva, A. C., & Souza, A. F. (2022). Spatial structure of the Caatinga woody flora: Abundance patterns have environmental, Pleistocene, and indigenous drivers. Anais Da Academia Brasileira de Ciências, 94(suppl 3), e20211019. 10.1590/0001-3765202220211019

57. Silva, J. L. S. E., Cruz-Neto, O., Peres, C. A., Tabarelli, M., & Lopes, A. V. (2019). Climate change will reduce suitable Caatinga dry forest habitat for endemic plants with disproportionate impacts on specialized reproductive strategies. PLOS ONE, 14(5), e0217028. 10.1371/journal.pone.0217028

58. Siyum, Z. G. (2020). Tropical dry forest dynamics in the context of climate change: Syntheses of drivers, gaps, and management perspectives. Ecological Processes, 9(1), 25. 10.1186/s13717-020-00229-6

59. Smith-Martin, C. M., Muscarella, R., Hammond, W. M., Jansen, S., Brodribb, T. J., Choat, B., Johnson, D. M., Vargas-G, G., & Uriarte, M. (2023). Hydraulic variability of tropical forests is largely independent of water availability. Ecology Letters, 26(11), 1829–1839. 10.1111/ele.14314

60. Souza, C. R. de, Santos, A. B. M., Maia, V. A., Paula, G. G. P. de, Fagundes, N. C. A., Coelho, P. A., Santos, P. F., Morel, J. D., Garcia, P. O., & Santos, R. M. dos. (2021). Seasonally dry tropical forest temporal patterns are marked by floristic stability and structural changes. CERNE, 27, e-102355. 10.1590/01047760202127012355

61. Stevens, J., Faralli, M., Wall, S., Stamford, J. D., & Lawson, T. (2021). Chapter 2 Stomatal Responses to Climate Change. In K. M. Becklin, J. K. Ward, & D. A. Way (Eds.), Photosynthesis, Respiration, and Climate Change (pp. 17–47). Springer International Publishing. 10.1007/978-3-030-64926-52

62. Terra, M. de C. N. S., Santos, R. M. dos, Prado Júnior, J. A. do, de Mello, J. M., Scolforo, J. R. S., Fontes, M. A. L., Schiavini, I., dos Reis, A. A., Bueno, I. T., Magnago, L. F. S., & ter Steege, H. (2018). Water availability drives gradients of tree diversity, structure and functional traits in the Atlantic–Cerrado–Caatinga transition, Brazil. Journal of Plant Ecology, 11(6), 803–814. 10.1093/jpe/rty017

63. Valavi, R., Elith, J., Lahoz-Monfort, J., & Guillera-Arroita, G. (2018). blockCV: An R package for generating spatially or environmentally separated folds for k-fold cross-validation of species distribution models. 10.1101/357798

64. Viana Santos, H., Borges De Lima, R., Figueiredo De Souza, R., Cardoso, D., Moonlight, P., Teixeira Silva, T., Pereira De Oliveira, C., Alves Júnior, F., Veenendaal, E., Paganucci De Queiroz, L., Rodrigues, P., Dos Santos, R., Sarkinen, T., De Paula, A., Barreto-Garcia, P., Pennington, T., & Phillips, O. (2023). Spatial distribution of aboveground biomass stock in tropical dry forest in Brazil. iForest – Biogeosciences and Forestry, 16(2), 116–126. 10.3832/ifor4104-016

65. Xie, H., Tian, S., Li, F., Sun, J., Hu, X., Bai, H., & Shao, C. (2025). Contrasting diurnal impacts of vapor pressure deficit on water use efficiency in two semiarid steppe ecosystems. Ecological Processes, 14(1), 68. 10.1186/s13717-025-00636-7

66. Zomer, R. J., Xu, J., & Trabucco, A. (2022). Version 3 of the Global Aridity Index and Potential Evapotranspiration Database. Scientific Data, 9(1), 409. 10.1038/s41597-022-01493-1

